# Understanding Research Development Needs in Capacity Building for NCD Research: The Case of Thailand

**DOI:** 10.1101/231498

**Authors:** N Singha-Dong, A Bigelow, P Furspan, B Rajataramya, A Villarruel, D Barton, K Potempa

## Abstract

Chronic non-communicable diseases (NCD) are the most significant causes of death globally. In Thailand, NCDs have increased 10.4% and 11%, respectively, since 2002. Thus, there is a compelling need in Thailand to enhance the capacity for research aimed at improving both NCD prevention and care. A survey was conducted of current multi-disciplinary doctorally prepared faculty (n=115) to determine perceived NCD research training needs. The results of this survey showed that the greatest exposure to NCD was in clinical practice, followed by teaching NCD content, and then research. Few researchers published their findings in journals. All responders reported needing significant support in research design, methods and statistical analysis procedures. These results were used to guide the development of a post-doctoral training program for NCD research in Thailand. After three years of the training program, we found that trainee applicants’ preferences and choices were aligned with the original survey-based planning.

## Introduction

Chronic non-communicable diseases (NCD) are the most significant causes of death globally. The WHO predicts that nearly half of the 89 million NCD-related premature deaths projected world-wide for the coming years will occur in the South East Asian region.^1,2^

Similar to other South-East Asian countries, chronic NCDs represent the largest cause of mortality in the Thai population.^3-5^ There also is strong evidence that the burden of chronic diseases is growing rapidly.^4,6-10^

Given the dramatic increase in morbidity and mortality in Thailand related to NCDs, a tailored research capacity building program was needed to provide a healthcare workforce poised to dramatically impact NCDs in Thailand.

We began our capacity building partnership for research with Thailand in 2010 with a D-71 planning grant. One of the first efforts was to better understand the training needs of PhD prepared nurses and other health professionals to determine the needs for training in NCD research. We then created the training plans for both short term and long term post-doctoral fellows in our NIH/Fogarty D43 research training program. Now three years into the program, we present here a brief report of whether our initial survey driven understanding of need was on target with the ongoing needs of the trainees.

For the formal survey the population of interest was all research trained faculty employed by the Praboromarajchanok Institute for Health Workforce Development [PIHWD], Ministry of Public Health in Thailand, which is a large training institution in Thailand for public health related personnel. The PIHWD employs a total of 200 doctoral faculty members at 38 colleges across Thailand in the disciplines of public health, nursing, pharmacy, medicine, and dentistry.

## Methods

### The Survey

We developed a survey questionnaire with 45 self-report items. The five-part questionnaire was developed to identify professional and educational background, past research and NCD experiences, training needs in research capacity as well as NCD specific training; the level and type of training needed; desired training locations; preferred delivery methods (e.g. course, shared experiences, distance learning); and a self-evaluation of communication skills in English. The survey had a variety of question types including open-ended questions, and questions with Likert-type responses. Text space was available for additional comment. Content validity was established through the review of the survey questionnaire by peers and subsequently an expert panel comprised of NCD researchers. Pilot testing with individuals similar to potential study participants identified issues with question wording, layout, understanding, or respondent reaction. Consequently, procedures, questionnaires, and documentation were modified and finalized according to the aforementioned evaluation.

### Procedure

A total of 200 doctoral prepared faculty members were invited to participate in the survey. Faculty who agreed to participate in the study signed the study consent form and responded to the paper-pencil questionnaire. The survey protocol was reviewed and approved by the Institutional Review Board of human subjects of the MOPH in Thailand.

## Results

After being reviewed for completeness, returned questionnaires were coded. Ten percent of the questionnaires were coded by two different individuals to ensure accuracy. Data were entered and then analyzed using descriptive and comparative statistics.

Of the 200 surveys distributed, 115 (57.5%) were completed and returned. Ninety-six of the respondents (83.4%) were female and 19 (16.5%) were male. The vast majority (*n*=96, 83.5%) of participants had a background in nursing. The average age of study participants was 45 years (*Range* 31-57 years, *SD* 5.35). In terms of doctoral discipline, 42 participants (36.5%) had a degree in nursing, 39 participants (33.9%) obtained a doctoral degree in education, 21 (18.2%) had a degree in other health and non-health related fields. The number of year post completion of doctoral degree varied from 0 to 12 years with an average of 3.42 years (*SD* 3.13).

Fifty-seven participants (50%) completed a doctoral program offered in English. Of the 80 who attended schools in Thailand, 22 (19.3%) participated in an English or international program.

### NCD Experience

The majority of participants (*n*=89, 77.39%) reported that they had had experience in NCD. These experiences varied, but included teaching, clinical supervision, training, and conducting research. Participants reported their NCD experience in terms of research, teaching, training, and other experience (52.17%, 48.70%, 22.61 %, and 10.43%, respectively). Fewer than half of the 64 participants who stated that they had conducted NCD studies reported that they had published their work. The average number of NCD studies and publications were 1.59 (*SD* = 0.49) and 1.38 (*SD* = 0.49), respectively.

### Overall Research Experience

In terms of research design, faculty reported conducting research and statistical methods using survey (n=92, 80%), correlational (n=68, 59%), comparative (n=56, 48.7%), experimental (n=32, 27.8%), and qualitative designs (*n*= 15, 13.4%). Most faculty (*n*=108, 93.9%) planned to conduct research in the future.

When asked about availability of time for conducting research, 46 (44%) responded often; 18 (15.7%) responded regularly; and 10 (8.7%) reported that they sometimes have the time to conduct research. A third (38 participants, 33%) mentioned that they rarely have time to do research and 2 participants (1.7%) reported no time for research in their current position.

### Post-Doctoral training needs as expressed in the survey

All participants expressed a need for building research capacity. Study participants identified training needs in the areas of health disparity (29%), diabetes (22.2%), cancer (17%), CVD (15%), hypertension (11.1%), and lung health (5.2%).

In terms of research skills, participants identified their training needs in all categories. From a total score of 5 for each skill, 5 indicating a strongest perceived need for training, average perceptions of research training scores are shown in Table 1.

**Table 1.**
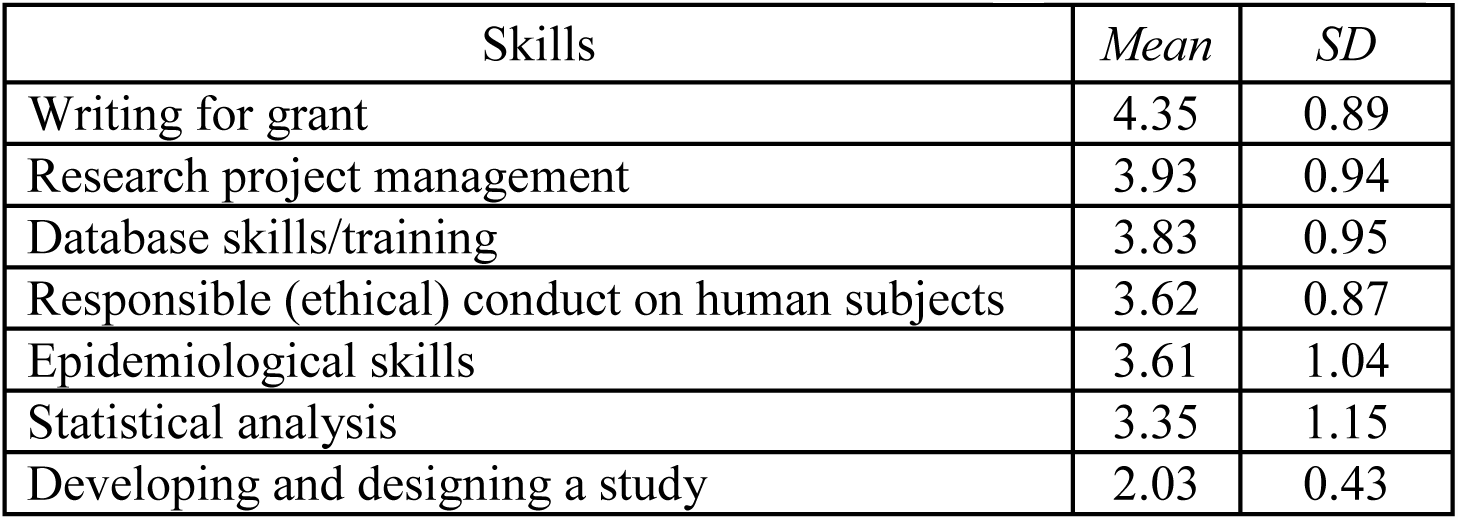
Means and Standard Deviations of training needs in research capacity skills

## Discussion

Given the formal survey results we designed a program that would fit our then understanding of the information provided. Overall we found that faculty members, regardless of disciplinary background, had various levels of NCD experience as well as research background and training. Thus, our design choice was to offer both long-term and short-term training options. Now, some three years after the start of our formal training programs, we find that the focus areas of our applicants, i.e., cancer, diabetes, hypertension, mental health disorders, prevention and chronic disease consequence of HIV, have corresponded well to the survey results and our understanding of the NCD situation in Thailand. Because the long-term, two-year post-docs were required to engage in every aspect of the mentor’s research activities they have received informal, but rigorous, training in project management, IRB applications, grant writing and manuscript/presentation preparation – top priority areas identified in the survey. The courses chosen by the long- and short-term trainees (Table 2) from the wide array available to them covered most of the areas indicated in the survey results (Table 1). The post-docs found that the courses were all quite advanced from those taken during their PhD training. Mastery of English in speaking and writing, a reported issue, was addressed through trainee engagement in seminars and oral presentations, and the preparation of manuscripts.

**Table 2.** Courses taken by long- and short-term trainees

|  |
| --- |
| Analysis, Statistics, and Survey Design<br>• Complex Systems Modeling for Public Health Research<br>• Summer Program in Quantitative Methods of Social Research<br>• Mixed Method Research Design and Data Collection<br>• An Introduction to Multilevel Analysis in Public Health<br>• Nursing Research: Survey Design and Analysis Using Mixed Methods<br>• Process tracing in qualitative and mixed methods research<br>• Structural equation models and latent variables: an introduction<br>• Structural equation modeling with STATA<br>• R statistics computing environment |

Another finding of the survey was perceived issues related to ‘time dedicated to research’. Through meetings with research directors/deans we learned that as expected, there is variability in the institutional priority given to research even in reputed research intensive institutions. A major effort is underway to advance institutional research capacity that supports faculty dedicated research effort, infrastructure supports, and productivity assessments. The indication of this need in the survey was helpful, if not instrumental, in bringing the project team leaders and the institutional research leaders together in various formats to further understand and design strategies for advancing institutional research capacity.

## Conclusion

We found that a formal survey provided reliable results in designing our approach to building NCD research capacity in Thailand. The benefits to this approach are two-fold; providing evidence to support capacity building design that reduces trial and error and, providing information that administrative stake holders find valuable in discussions to make institutional changes necessary to advance research capacity.

